# MicroRNA-378 suppressed osteogenesis of mesenchymal stem cells and impaired bone formation via inactivating Wnt/β-catenin signaling

**DOI:** 10.1101/699355

**Authors:** Lu Feng, Jin-fang Zhang, Liu Shi, Zheng-meng Yang, Tian-yi Wu, Hai-xing Wang, Wei-ping Lin, Ying-fei Lu, Jessica Hiu Tung Lo, Da-hai Zhu, Gang Li

## Abstract

MicroRNAs (miRNAs) have been reported to serve as silencers to repress gene expression at post-transcriptional level. Multiple miRNAs have been demonstrated to play important roles in osteogenesis. MiR-378, a conserved miRNA, was reported to mediate bone metabolism and influence bone development, but the detailed function and underlying mechanism remain obscure. In this study, the miR-378 transgenic (TG) mouse was developed to study the role of miR-378 in osteogenic differentiation as well as bone formation. The abnormal bone tissues and impaired bone quality were displayed in the miR-378 TG mice, and a delayed healing effect was observed during bone fracture of the miR-378 TG mice. The osteogenic differentiation of MSCs derived from this TG mouse was also inhibited. We also found that miR-378 mimics suppressed while anti-miR-378 promoted osteogenesis of human MSCs. Two Wnt family members Wnt6 and Wnt10a were identified as *bona fide* targets of miR-378, and their expression were decreased by this miRNA, which eventually induced the inactivation of Wnt/β-catenin signaling. Finally, the sh-miR-378 modified MSCs were locally injected into the fracture sites in an established mouse fracture model. The results indicated that miR-378 inhibitor therapy could promote bone formation and stimulate healing process *in vivo*. In conclude, miR-378 suppressed osteogenesis and bone formation *via* inactivating Wnt/β-catenin signaling, suggesting miR-378 may be a potential therapeutic target for bone diseases.

## Introduction

Bone regeneration is very important for the recovery of some diseases including osteoporosis and bone fracture trauma. It is a multiple step and multiple gene involved complex process, including osteoblast-mediated bone formation and osteoclast-mediated bone resorption. During this process, mesenchymal stem cells (MSCs) gradually differentiate into osteoblasts, which produce a variety of extracellular matrix and thus induce the initiation of bone formation. Hence, improving the osteoblast differentiation of MSCs is crucial for the development of therapeutic strategy for bone diseases.

Over the past few years, microRNAs (miRNAs) have emerged as gene silencers to suppress gene expression at post-transcriptional level. Multiple miRNAs have been demonstrated to play important roles in biological activities including osteogenic differentiation and skeletal development. For example, miR-26a, miR-29b, miR-125b, miR-133, miR-135 and miR-196a, have been reported to be involved in osteogenesis.^1–5^ Our previous reports also demonstrated that miR-20a and 20b promoted whereas miR-637 suppressed osteoblast differentiation.^6–7^ Therefore, miRNAs have been considered as potential candidates to mediate osteogenic differentiation and may be acted as therapeutic targets for bone regeneration.

MiR-378, which is derived from a hairpin RNA precursor, has been reported to promote BMP2-induced osteoblast differentiation in C2C12 cells^8^ and attenuate high glucose-suppressed osteogenic differentiation in preosteoblastic cell line,^9^ demonstrating the promoting effect on osteoblast. More strangely, other group reported that miR-378 suppressed osteogenic differentiation of MC373-E1 cells.^10^ Therefore, the function of miR-378 in osteogenesis remains intriguing.

In this study, abnormal bone tissues and impaired bone quality were exhibited in the miR-378 transgenic (TG) mice; and a delayed bone formation and fracture healing was also found in this TG mice. Moreover, we found that miR-378 overexpression suppressed while its antagonists promoted osteogenic differentiation, suggesting miR-378 may play a significant role in osteoblast differentiation and bone regeneration. Furthermore, two Wnt family members Wnt6 and Wnt10a were identified as targets of miR-378, and their expression were decreased by miR-378, which eventually suppressed Wnt/β-catenin signaling. Therefore, miR-378 inhibited osteoblast differentiation and impaired bone formation, suggesting that it may be a potential therapeutic target for bone diseases.

## Results

### Abnormal bone tissues and impaired bone quality were observed in miR-378 TG mice

To investigate the function of miR-378 in bone formation, we compared the bone tissues from miR-378 TG mice and that from their wild-type (WT) mice. By digital radiography examination, the bone size and length of the femur (Figure 1A), tibia (Figure 1B), head (Supplementary Figure 1A for side view and Supplementary Figure 1B for vertical view), tail (Supplementary Figure 1C) and spine (Supplementary Figure 1D for front view and Supplementary Figure 1E for side view) were all decreased in the miR-378 TG mice. We further investigated the microarchitecture of femurs and tibia by micro-CT examination. The representative 3D images showed the lower bone mass in cortical and trabecular bone of femur (Figure 1C-1D) as well as that of tibia (Figure 1E-1F) in miR-378 TG mice. Furthermore, hematoxylin-eosin (H&E) staining of femurs showed that the thinner growth plate, less smooth edge of cortical bone and less paralleled alignment of osteocytes in TG mice (Figure 1G), and less trabecular bone mass around the metaphyseal region of femurs in miR-378 TG mice, suggesting more bone lost in cancellous bone (Figure 1H).

**Figure 1.**
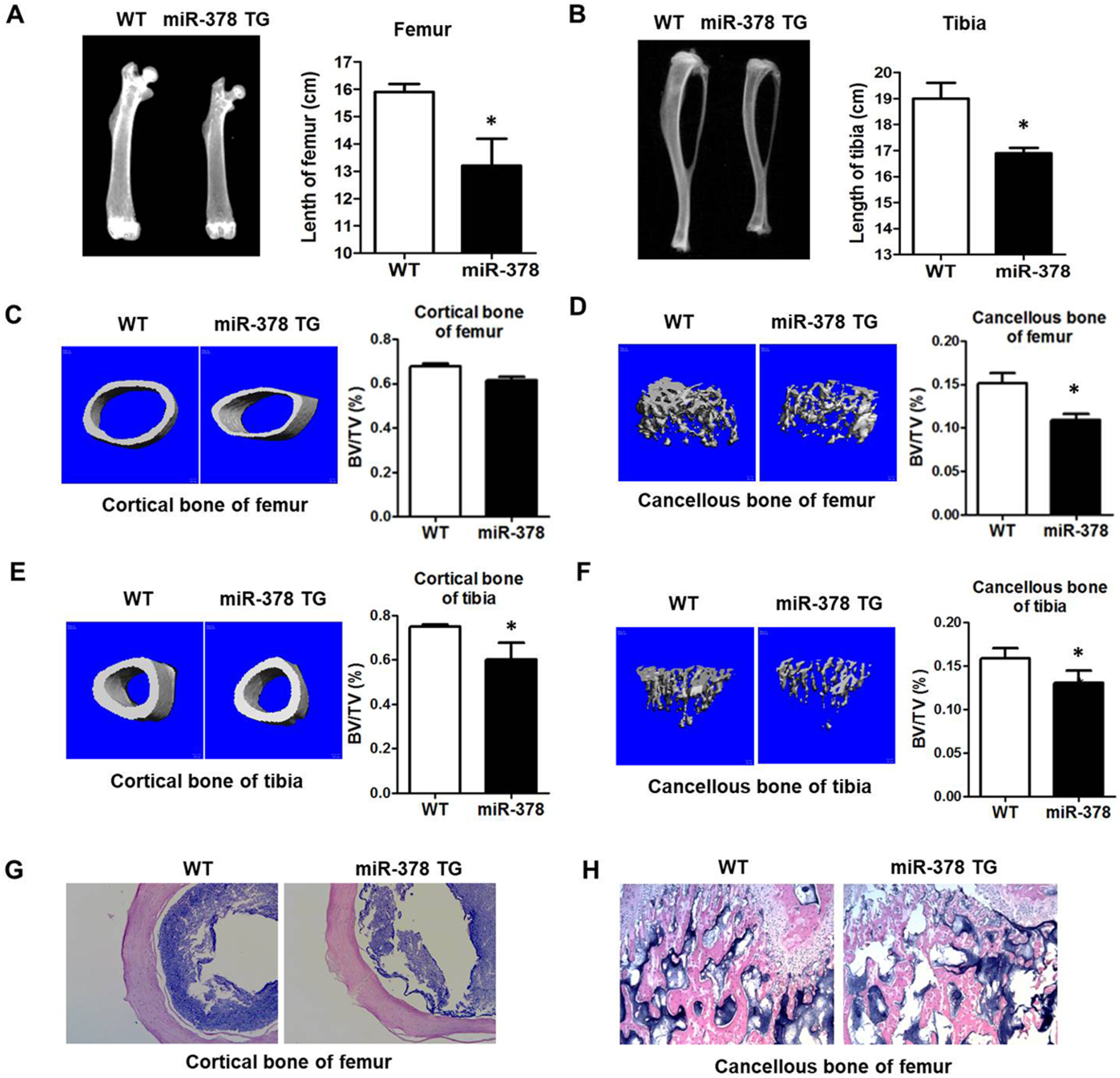
Abnormal bone tissues and impaired bone quality were observed in miR-378 TG mice. A-B, The bone phenotype of femurs (A) and tibia (B) of miR-378 TG mice and their wild-type (WT) mice were examined by digital radiography. **C-F**, Representative microarchitecture 3D images of cortical (C, E) and trabecular bone (D, F) of femur as well as tibia. Bone volume over total volume (BV/TV) of the cortical and trabecular bone all showed that the bone mass was significantly decreased in the miR-378 TG mice. **G**, H&E staining of transverse and coronal plane of femurs revealed thicker cortices, smoother edge of cortical bone and more paralleled alignment of osteocytes in the WT mice than those in the miR-378 TG group. **H**, H&E staining of sagittal plane of metaphyseal region of femurs revealed more bone lost in cancellous bone of miR-378 TG mice femur. (Age: 3 month; n = 10; *, P < 0.05).

### Bone fracture healing was delayed in miR-378 TG mice

Following the observed abnormal bone tissues in miR-378 TG mice, an established mouse femoral fracture model was performed to compare the healing process of bone fracture between miR-378 TG and WT mice. The healing process were assayed by X-ray examination, and the representative images showed that less callus formation was observed in the miR-378 TG group compared with the WT group during the healing process (Figure 2A). At week 4, the gap in the fracture sites disappeared in WT group while it also appeared in miR-378 TG group, indicating that the fracture healing process was delayed in miR-378 TG mice. The 3D reconstructed images of μCT also confirmed the lower bone mass in the TG mice at week 4 (Figure 2B); and the quantitative analysis showed that the TG group had significantly decreased BV/TV, suggesting less newly formed mineralized bone in TG group (Figure 2C). Furthermore, the three-point bending mechanical testing was performed and results showed that the miR-378 TG group had a significant decrease in ultimate load, E-modulus and energy to failure (Figure 2D). H&E staining of fracture healing position revealed that the fracture site of WT group was covered by more calluses and the bone remodeling was more vigorously compared with TG group (Figure 2E). Moreover, by immunohistochemistry staining, the decreased OCN and OPN expression were also observed in miR-378 group, suggesting an impaired effect of miR-378 on bone formation and remodeling (Figure 2E).

**Figure 2.**
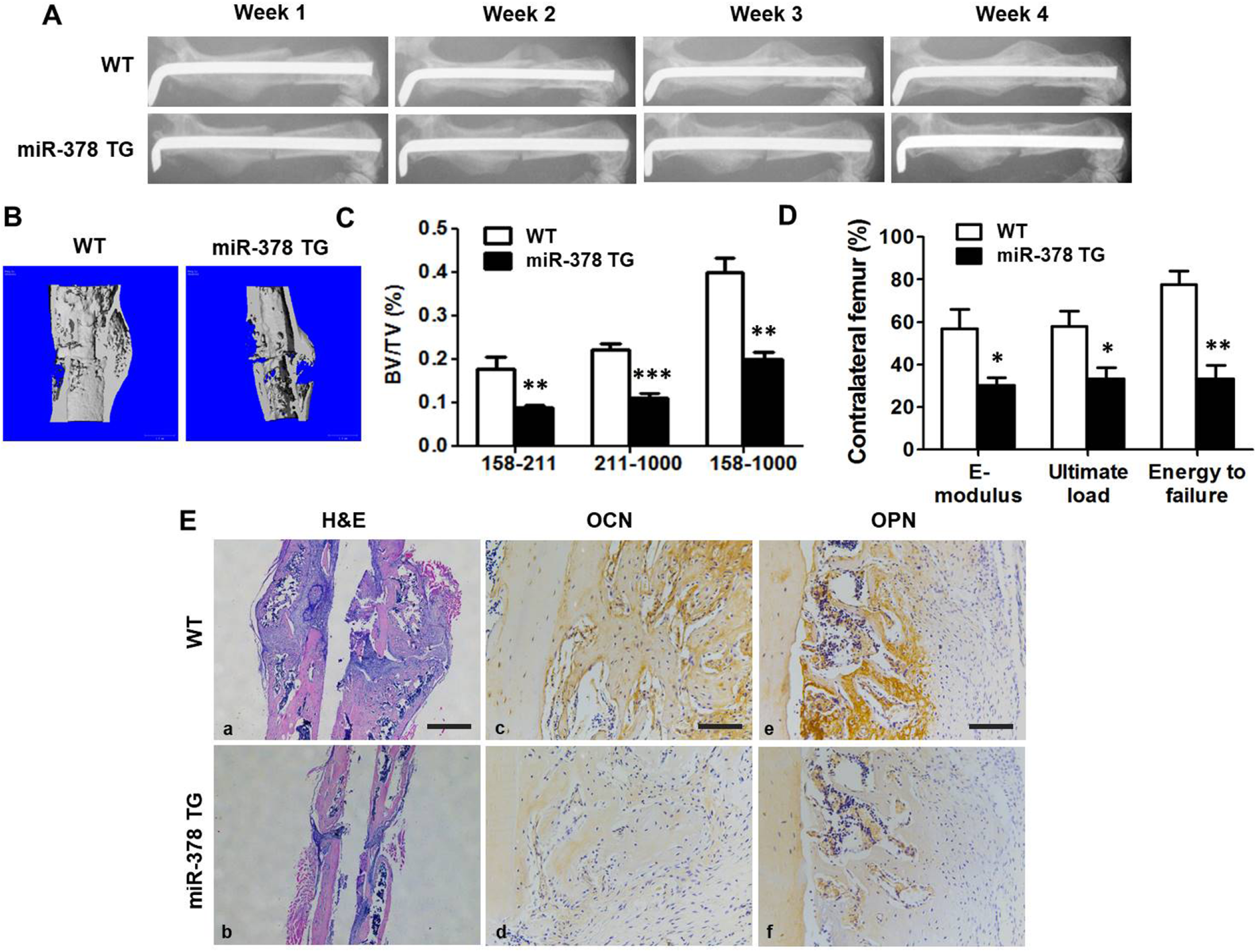
Bone fracture healing was impaired in miR-378 TG group. **A**, Representative images of X-ray radiography showed that less callus formation was found in the miR-378 TG group at week 4, and the gap in the fracture sites almost disappeared in WT group. **B**, Three-dimension μCT images of the mouse femur fracture zone were taken 4 weeks after surgery, and the result confirmed less continuous callus was found in miR-378 TG group. **C**, Micro-CT analysis was performed and highly mineralized and newly formed callus were reconstructed based on different threshold. The results showed that bone volume/total tissue volume (BV/TV) of total mineralized bone (threshold 158-1000), highly mineralized bone (threshold 211-1000) as well as new callus (threshold 158-211) in the WT group was much higher than that in the miR-378 TG groups. **D**, Three-point bending mechanical testing in the miR-378 TG group showed significant decrease of ultimate load, E-modulus, and energy to failure compared to WT group. **E**, (**a-b**), H&E staining revealed miR-378 TG group has fewer calluses covered on femur fracture site at week 4 post-surgery and the bone remodeling was less vigorous than WT group. Scale bar = 800 µm. (**c-f)**, Less Osx and OCN expression was in the miR-378 TG group by immunohistochemistry staining. Scale bar = 100 µm. (Age: 3 month; n = 10; *, P < 0.05; **, P < 0.01; ***, P < 0.001).

### MiR-378 suppressed osteogenic differentiation of MSCs

To analyze the involvement of miR-378 during osteogenesis, we next investigate the osteogenic potential of MSCs derived from miR-378 TG mouse. Under osteogenic inductive condition, alkaline phosphatase (ALP) activity, an early marker of osteogenesis, was lower in MSCs derived from miR-378 TG group when compared with that from WT group at day 3 (Figure 3A). And the TG group had fewer mineralized nodules compared to that of the WT group at day 14 (Figure 3B). Furthermore, the expression of osteogenic marker genes including Runt-related transcription factor 2 (Runx2), ALP, osteocalcin (OCN), osteopontin (OPN), osteoprotegerin (OPG), bone morphogenetic protein 2 (BMP2) and osterix (Osx) were significantly repressed in the TG group at day 7 (Figure 3C).

**Figure 3.**
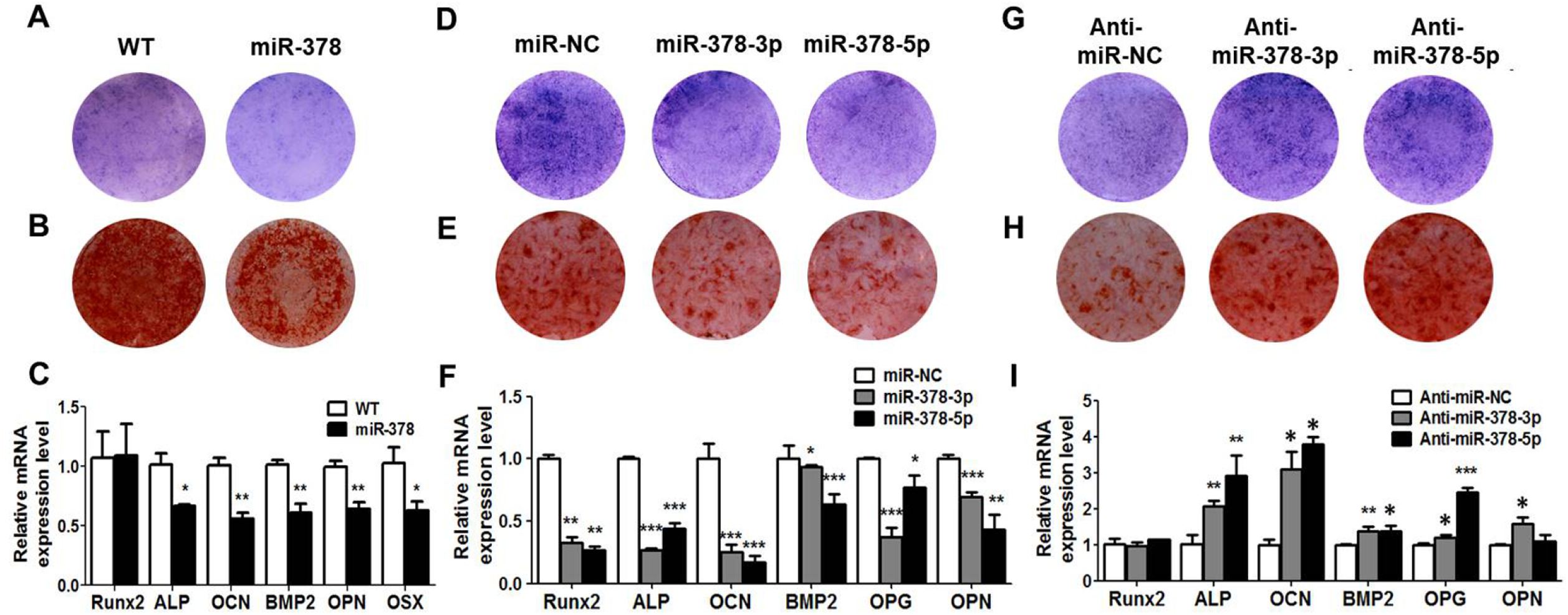
miR-378 suppressed osteogenesis of MSCs. A-C, Under osteogenic induction, ALP activity (A), calcified nodules (B) and osteogenic marker genes expression (C) were all suppressed in MSCs derived from miR-378 TG mice. **D-F**, ALP activity (D), calcified nodules formation (E) and osteogenic marker genes expression (F) were all decreased by miR-378-3p or miR-378-5p mimics. **G-I**, ALP activity (G), calcified nodules formation (H) and osteogenesis marker genes expression (I) were promoted by anti-miR-378-3p and anti-miR-378-5p. (n = 3; *, P < 0.05; **, P < 0.01; ***, P < 0.001).

To further identify the function of miR-378 in osteoblast differentiation, miR-378 mimics and antagonist were also introduced into human bone marrow MSCs under osteogenic inductive condition. ALP activities were suppressed by miR-378-3p and miR-378-5p mimics at day 3 (Figure 3D). And fewer calcium nodules were observed in miR-378-3p and miR-378-5p groups at day 14 (Figure 3E). The expression of osteogenic marker genes including Runx2, ALP, OCN, BMP2, OPN and Osx were significantly suppressed by miR-378-3p and miR-378-5p mimics (Figure 3F). On the other hand, when the antagonists of miR-378-5p and miR-378-3p were introduced, the ALP activity (Figure 3G), mineralized nodules (Figure 3H) and the osteogenic marker genes expression (Figure 3I) were obviously increased. All these data demonstrate that miR-378 may directly regulate the osteogenesis of MSCs, thereby affecting bone development.

### Wnt6 and Wnt10a were real targets for miR-378 in MSCs

MiRNAs function as regulators in multiple biological activities through directly targeting protein-coding genes. In terms of the suppressive effect of miR-378 on osteogenesis, we tried to characterize the candidate target genes of this miRNA. Among the candidates predicted by bioinformatics analyses, the Wnt family members Wnt6 and Wnt10a were found to be the most promising candidates for miR-378-3p and miR-378-5p, respectively. Their predicted binding sites were shown in Figure 4A and Figure 4C. To validate their direct interaction, the binding and mutated sites into the 3’UTR of Wnt6 and Wnt10a were inserted into the pmiR-GLO vector to generate the luciferase reporter vectors. Co-transfection of miR-378 isoforms with the two luciferase reporters was performed, and it was showed that miR-378-5p and miR-378-3p dramatically suppressed the luciferase activity of these luciferase reporters of Wnt6 and Wnt10a and mutations on their binding sites successfully abolished the suppressive effects (Figure 4B and Figure 4D). With miR-378-3p and miR-378-5p transfection, the expression of Wnt10a and Wnt6 were both significantly reduced in human BMSCs at the mRNA and protein levels (Figure 4E and Figure 4G). On the contrary, both anti-miR-378-3p and anti-miR-378-5p promoted the expression of Wnt10a and Wnt6 at miRNA and protein levels, respectively (Figure 4F & Figure 4H).

**Figure 4.**
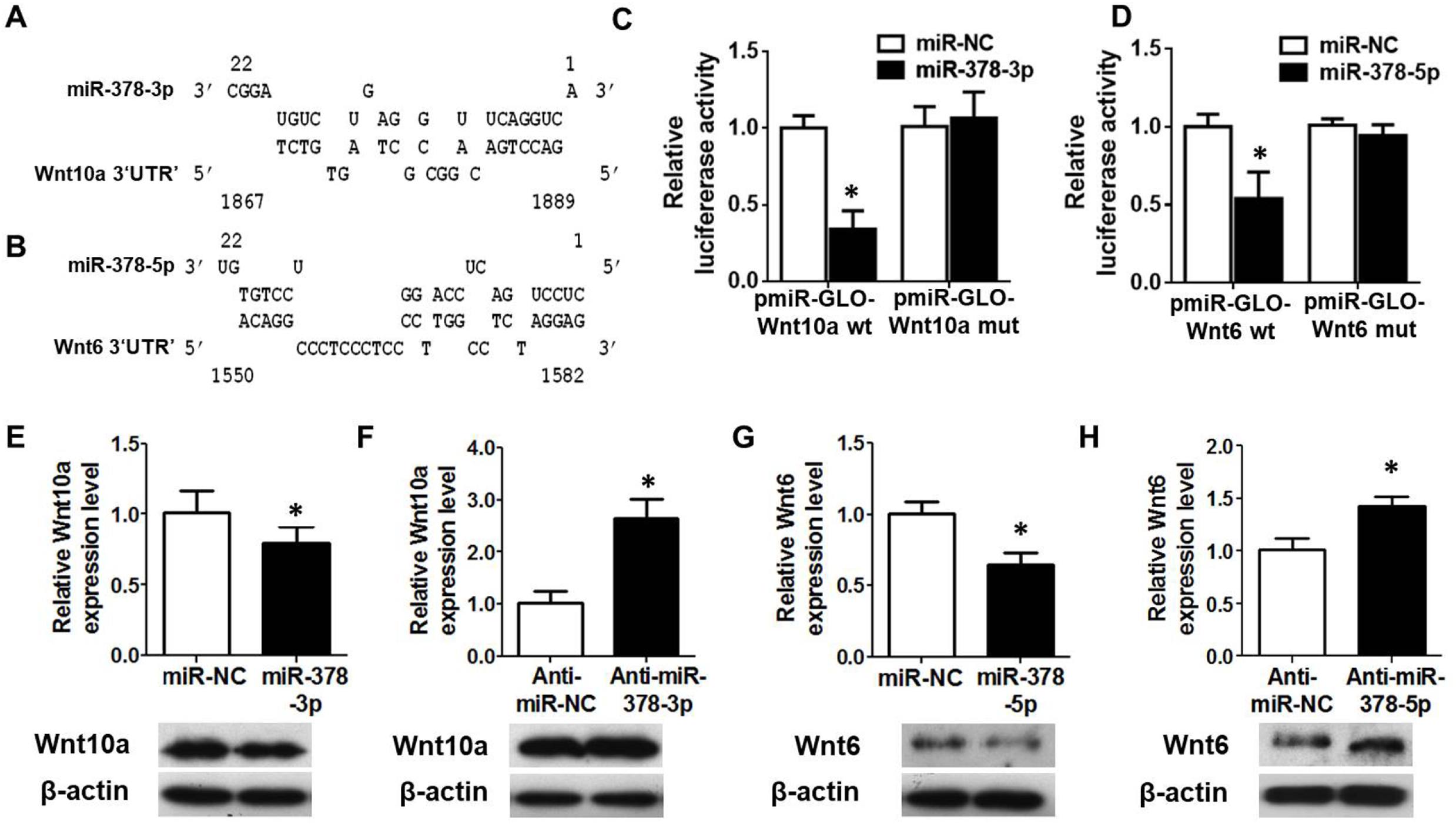
Wnt10a and Wnt6 were *bona fide* targets for miR-378-3p and miR-378-5p, respectively. A-B, Schematic diagrams of the interaction between miR-378-3p and miR-378-5p transcripts and their direct target genes Wnt10a (A) and Wnt6 (B). **C-D**, MiR-378-3p or miR-378-5p were co-transfected with the Wnt10a or Wnt6 luciferase reporter into HEK293 cells, and the luciferase activity were measured. **E-F**, the expression of Wnt10a was suppressed by miR-378-3p mimics (E) while promoted by its antagonist (F) at mRNA and protein levels. **G-H**, the expression of Wnt6 was reduced by miR-378-5p mimics (G) while enhanced by its antagonist (H) at mRNA and protein levels. (n = 3; *P < 0.05; **P < 0.01; ***P < 0.001).

### Wnt/β-catenin signaling was involved in miR-378-mediated osteogenesis

Considering that both Wnt6 and Wnt10a were the members of Wnt family, we next investigated whether Wnt/β-catenin signaling could be involved in the miR-378 mediated osteogenesis. The Wnt signaling reporter TOPflash, which contains three binding sites for TCF and β-catenin, was introduced. The results showed that both miR-378-3p and miR-378-5p reduced while their inhibitors activated the luciferase activities (Figure 5A-5B). Consistent with these results, we also found that miR-378-3p and miR-378-5p suppressed while their inhibitors promoted β-catenin expression at mRNA and protein levels (Figure 5C-5D). Furthermore, several downstream transcriptional targets such as c-Myc, CD44 and cyclin D1 were downregulated by ectopic expression of miR-378 isoforms (Figure 5E), while upregulated by anti-miR-378(Figure 5F). Expression level of several other Wnt family members activating Wnt/β-catenin signaling pathway were also downregulated in MSCs isolated from miR-378 TG mice, which revealed a broader influence of miR-378 overexpression on Wnt/β-catenin signaling mediated osteogenesis (Supplementary Figure 2).

**Figure 5.**
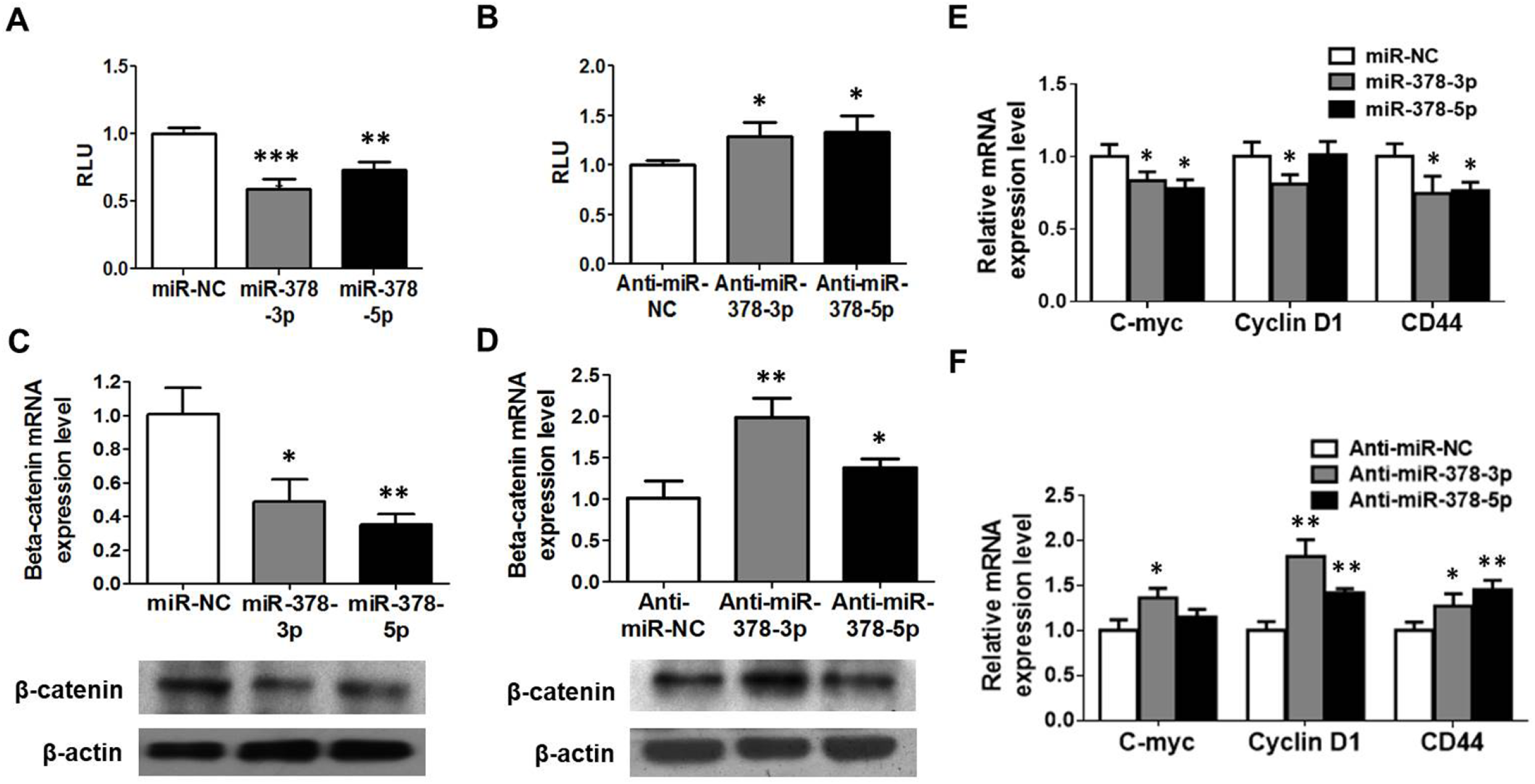
Wnt/β-catenin signaling was suppressed in miR-378 mediated osteogenesis. **A-B**, miR-378 mimics suppressed while its antagonist promoted the TOPflash luciferase activity. **C-F**, The expression of β-catenin was evaluated with miR-378 mimics or the antagonists treatment at mRNA (C, E) and protein (D, F) levels. **G-H**, miR-378 mimics (G) suppressed while its antagonists (H) activated several downstream target genes of β-catenin. (n = 3; *, P < 0.05; **, P < 0.01; ***, P < 0.001).

To further confirm the involvement of Wnt/β-catenin signaling *in vivo*, the MSCs derived from miR-378 TG mouse was used for investigation. Downregulated Wnt6 (Figure 6A), Wnt10a (Figure 6B) and β-catenin (Figure 6C) were observed in the MSCs derived from miR-378 TG group. Consistent with the mRNA results, their protein expression was all decreased in MSCs from TG group (Figure 6D-6E). And the Wnt/β-catenin signaling downstream target genes such as c-Myc, CD44 and cyclin D1 were significantly downregulated in MSCs derived from miR-378 TG mouse (Figure 6F). All these data suggest that Wnt/β-catenin signaling could be involved in miR-378 mediated osteogenesis.

**Figure 6.**
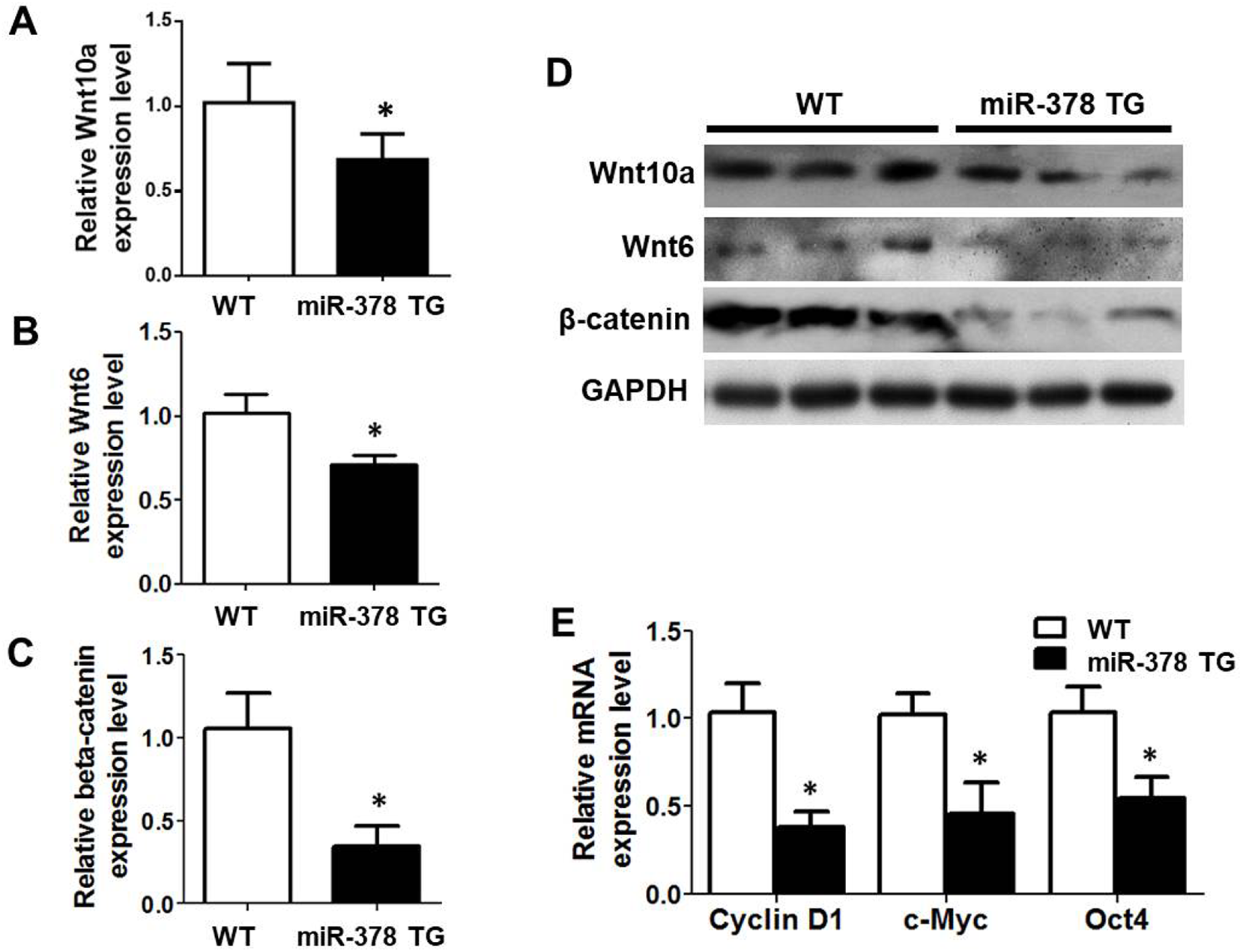
Wnt/β-catenin signaling was suppressed in miR-378 TG mice. **A-C**, expression of Wnt10a (A), Wnt6 (B) and β-catenin (C) was decreased in the MSCs derived from miR-378 TG mice at mRNA level. **D**, The expression of Wnt10a, Wnt6 and β-catenin was suppressed in the MSCs isolated from miR-378 TG mice. **E**, several downstream target genes of β-catenin were downregulated in the MSCs derived from miR-378 TG mice. (n = 3; *, P < 0.05).

### Sh-miR-378 modified MSCs accelerated bone fracture healing *in vivo*

To further investigate the *in vivo* effect of miR-378 on bone formation, a mouse femoral fracture model was used in this study. The lentivirus particles were produced and mouse MSCs were infected with these lentiviruses. The sh-miR-378- and sh-NC-infected MSCs were locally injected into the fracture sites at day 3 post-surgery and X-ray examination were taken every week. The results showed that the gap in the fracture sites nearly disappeared at week 4 in sh-miR-378 group compared with control group (Figure 7A). Further micro-CT analyses showed more newly mineralized calluses in the sh-miR-378 group (Figure 7B). Besides, the value of bone volume/total tissue volume (BV/TV) was calculated and the results exhibited significant enhancement of newly formed mineralized bone (threshold 211-1000) and total mineralized bone (threshold 158-1000), indicating more newly formed mineralized bone in sh-miR-378 group (Figure 7C). To further evaluate the quality of fracture healing, mechanical testing was performed at week 4 and the results showed that the E-modulus, ultimate load and energy to failure were all significantly increased in sh-miR-378 group (Figure 7D). The sh-miR-378 and sh-NC-infected MSCs could express the green fluorescent protein (GFP) (Supplementary Figure 3A), and the GFP fluorescence of the slides confirmed the localization of these infected MSCs at the fracture callus (Supplementary Figure 3B). The HE and IHC staining were performed to evaluate the histological properties of newly mineralized-bone tissues. As shown in Figure 7E, the fracture site of sh-miR-378 group was covered by more calluses and the bone remodeling was more vigorously compared with control group at week 4 post-surgery. Moreover, the increased OCN and OPN expression were also observed in sh-miR-378 group (Figure 7E), which suggests a better effect of sh-miR-378 on bone formation and remodeling.

**Figure 7.**
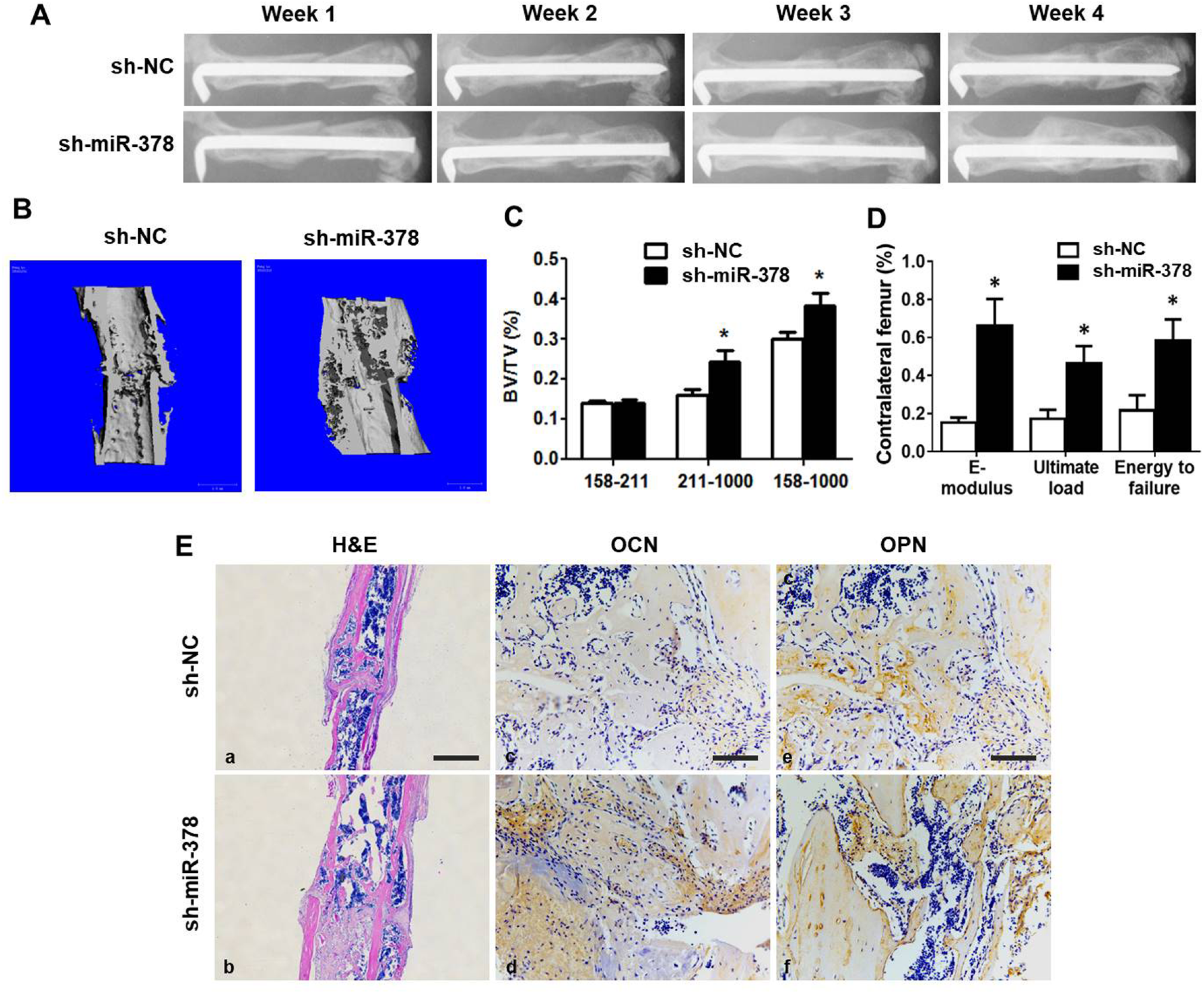
sh-miR-378 mediated MSCs accelerated bone fracture healing in mouse. A, X-ray radiography was taken during the course of fracture healing. Representative images showed that the fractured site was covered by more callus and bone remodeling was more vigorous in sh-miR-378 group. **B**, Three-dimension μCT images showed a better fracture healing effect condition and more callus in sh-miR-378 group. **C**, The statistical diagram of BV/TV calculated from micro-CT results was displayed. **D**, Three-point bending mechanical testing at week 4 post surgery was performed. The statistical data (B) were displayed. **E, (a-b)**, H&E staining revealed more callus covered in femur fracture site of sh-miR-378 group at week 4. Scale bar = 800 µm. **(c-f)**, More Osx and OCN expressed in the new bone zone in sh-miR-378 treatment group by immunohistochemistry staining. Scale bar = 100 µm. (Age: 3 months; n = 10; *, P < 0.05; **, P < 0.01; ***, P < 0.001).

## Discussion

With the aging of global population, osteoporosis has been becoming a serious public health problem, which is characterized by the decreased bone density and subdued strength, eventually leading to an increased risk of fracture. In the present study, miR-378 was found to suppress osteogenic differentiation *in vitro* and impair bone formation and fracture healing process as well *in vivo*, indicating that miR-378 may be a potential therapeutic target for bone diseases.

As a conserved miRNA, miR-378 is located in the first intron of peroxisome proliferator-activated receptor γ coactivator-1β (PGC-1β),^11^ and it is widely expressed in many tissues including adipose, skeletal muscles, myocardium, etc. MiR-378 was originally identified as an oncogene promoting VEGF expression in human nasopharyngeal carcinoma,^12^ now it has been proved to participate in a variety of biological processes, such as cancer metastasis and differentiation.^13–14^ MiR-378 was strongly upregulated during adipogenic differentiation and positively regulated adipogenesis.^15–17^ As for osteogenesis, some studies reported that miR-378 promoted osteogenic differentiation,^8^ while other study demonstrated that miR-378 suppressed osteogenic differentiation.^10^ Therefore, the exact function of miR-378 on osteogenesis remains unclear. Recently, Zhu’s group reported that skeletal muscle mass was significantly reduced in transgenic mice globally overexpressing miR-378 (TG) compared with that in the WT mice.^18^ In the present study, the abnormal bone tissues and impaired bone quality were observed in the miR-378 TG mouse, and moreover the bone fracture healing was delayed in the femoral fracture model of this transgenic mouse. Our study firstly indicated the *in vivo* activity of miR-378 in regulation bone development by using this transgenic animal model.

The *in vitro* effect of miR-378 on MSC osteogenesis was further examined in this study. The MTT assay was firstly performed to compare the proliferation activity of bone marrow MSCs from WT and miR-378 TG mice, the result concluded that miR-378 could not alter MSC proliferation (Supplementary Figure 4E). Under the osteogenic inductive conditions, MSCs derived from miR-378 TG mice showed weak potential of osteogenic differentiation compared with that from WT mice. Furthermore, we also found that miR-378 mimics suppressed while their inhibitors could promote osteogenic differentiation of human MSCs. All these provide strong support for the impaired bone formation in miR-378 TG mice. Consistent with our study, miR-378 inhibited osteogenesis of mouse osteoblast cell line MC3T3-E1 cells.^10^ miR-378 secreted by osteoclast was also discovered to be increased in exosomes of patients with bone metastases compared to healthy controls and the expression level was correlated with bone metastasis burden.^19^ Based on the previous reports and our results, miR-378 may be a negative regulator of osteogenesis and bone regeneration.

As for the molecular mechanism of miR-378, it has been reported that miR-378 mediated metabolic homeostasis in skeletal muscle via Akt1/FoxO1/PEPCK pathway. IGF1R signaling pathway was also reported to be involved in miR-378 mediated muscle regeneration. Wnt/β-catenin signaling was critical for normal bone and tooth formation and development.^20^ This pathway is essential for multiple biological activities including osteogenesis. Various studies have demonstrated that miR-378 could regulate Wnt/β-catenin signaling, i.e. miR-378 could increase neural stem cell differentiation through Wnt/β-catenin signaling;^21^ A colon cancer study also revealed that miR-378 attenuates malignant phenotypes of colon cancer cells *via* suppressing Wnt/β-catenin pathway.^22^ More importantly, miR-378a-3p could suppress Wnt/β-catenin signaling in hepatic stellate cells *via* targeting Wnt10a.^23^ In the current study, two Wnt family members, Wnt6 and Wnt10a were identified as the targets of miR-378, and overexpressed miR-378 could suppress their expression, thus resulted in inactivating Wnt/β-catenin signaling. As members of Wnt gene family, Wnt10a could induce MSCs osteoblastogenesis by activating and stabilizing the downstream β-catenin expression and inducing Wnt/β-catenin signaling;^24^ and Wnt6 could act synergistically with BMP9 to induce Wnt/β-catenin signaling as well as MSC osteogenic differentiation.^25^ Moreover, Wnt6 also promoted Runx2 promoter activity directly and stimulated osteogenesis.^26^ In *in vivo* studies, Wnt6 and Wnt10a were also revealed to be pivotal characters in bone development. For example, Wnt6 is expressed during long bone development.^27^ While the expression level of Wnt10a was also revealed downregulated in Runx2 knockout mice,^28^ as well as bone marrow MSCs isolated from ovariectomy induced osteoporosis mice.^29^ These data further supported that Wnt6 and Wnt10 is in bone metabolism and downregulated Wnt6 and Wnt10a was highly related to bone disorder disease. Taken together, the previous and our research results all indicated that miR-378 directly suppressed Wnt6 and Wnt10a mRNA expression, and hence represses Wnt/β-catenin signaling as well as osteogenic differentiation of MSCs.

To further investigate the *in vivo* therapeutic effect of miR-378, sh-miR-378-modified MSCs were applied to an established mouse femoral fracture model for bone fracture treatment. Our results demonstrated that local administration of miR-378 inhibitor modified cells promoted bone formation and improved mechanical properties of fractured femur. What’s more, the micro-CT examination showed a significant increase of newly formed callus and total mineralized bone volume in sh-miR-378 group. Furthermore, more vigorous bone formation and bone remodeling was observed in sh-miR-378 group by histological analyses. Therefore, these results suggest an accelerated effect of sh-miR-378 on fracture healing *in vivo*.

In summary, our data demonstrated that miR-378 could impair the bone formation *in vivo* and suppress the osteogenesis *in vitro*. Two Wnt family members, Wnt6 and Wnt10a, were identified as novel targets of this miRNA. MiR-378 led to the repression of the two targets, which eventually inactivated Wnt/β-catenin pathway and hence suppressed osteogenesis. Therefore, mR-378 may be a potential novel therapeutic target, and the knowledge gained from this study will provide insight for developing a new cell therapy strategy to bone diseases.

## Material & methods

### Plasmid generation and cell transfection

The miR-378 TG mice was developed and kindly provided by Prof. Dahai Zhu (Chinese Academy of Medical Sciences).^18^ The genotyping characterization of miR-378 TG mice was performed following the protocol from Zhu. *et al*’s research group and the result was described in Supplementary Figure 5. MiR-378 mimics and miR-378 inhibitors were purchased from GenePharma Company (Shanghai, China). Human Wnt10a and Wnt6 3’-UTR sequence were subcloned into pmiR-GLO vector. All the cDNA sequences were obtained by database searching (NCBI: http://www.ncbi.nlm.nih.gov/). Sh-miR-378 lentivirus plasmid was designed and constructed by GenePharma Company (Shanghai, China). MiRNA mimics and inhibitors as well as DNA plasmids were transfected using transfection reagent Lipofectamine 3000 (Invitrogen, USA) following the manufacturer’s instruction.

### Cell culture and osteo-induction

The human embryonic kidney 293 (HEK293) cells was purchased from ATCC and was cultured in Dulbecco’s modified Eagle medium (DMEM) supplemented with 10% heat-inactivated fetal bovine serum (FBS) and 1% penicillin/streptomycin/neomycin (PSN). The human bone marrow-derived mesenchymal stem cells (MSCs) were isolated from bone marrow, which aspirated from healthy donors. Mouse MSCs were isolated from bone marrow of 3-month-old miR-378 TG mouse, the WT mice at the same age was served as a negative control. Both human and mouse bone marrow was flushed out into a 10-cm cell culture dish using Minimum Essential Medium Alpha (MEMα) plus 10% heat-inactivated FBS and 1% PSN. The culture was kept in a humidified 5% CO2 incubator at 37°C for 72h, when non-adherent cells were removed by changing the medium. MSCs were characterized using flow cytometry for phenotypic markers including MSC positive markers CD44-FITC, CD90-PE and negative markers CD31-FITC and CD45-FITC following the previous protocol^6^ (Supplementary Figure 4A-4D) and the stocked in liquid nitrogen for future use.^30^ To initiate osteogenic differentiation, the MSCs were seeded in a 12-well plate and grow up to 80% confluence, and 10 nM dexamethasone (Sigma-Aldrich, USA), 50 μg/ml ascorbic acid (Sigma-Aldrich, USA), and 10 mM glycerol 2-phosphate (Sigma-Aldrich, USA) were added into the culture medium. The differentiation medium was replaced every 3 days.

### ALP activity and alizarin red S staining

Mouse or human MSCs were seeded in 24 well plates at a density of 2× 10^5^ cells per well and osteogenic differentiation was induced when cells reach 80% confluence. For the alizarin red S staining, the mouse or human MSCs were washed with PBS and fixed with 70% ethanol for 30 min. The MSCs were then stained with 2% alizarin red S staining solution for 10 min. The stained calcified nodules were scanned using Epson launches Perfection V850 (Seiko Epson, Japan). The ALP activity was measured and analyzed following the published protocol.^6^

### Establishment of stable cell lines

Sh-miR-378 mediated mouse MSCs were generated using lentivirus-mediated gene delivery system as previously described.^6^ And the supernatant medium containing lentivirus particles were purchased from GenePhama (Shanghai, China). The mouse MSCs were infected with the lentiviral particles by the addition of hexadimethrine bromide (Sigma-Aldrich, USA). Lentivirus infected mouse MSCs were selected by G418 (Sigma-Aldrich, USA) at 500μg/ml. After antibiotics selection for around 7 days, remained cultured cells were collected and knockdown of miR-378 was confirmed by qRT-PCR examination.

### RNA extraction and qRT-PCR examination

Total RNA of the cultured cells were harvested with RNAiso Plus reagent (TaKaRa, Japan) following the manufacture’s instruction. After RNA extraction, cDNA were reversely transcribed from RNA samples by PrimeScript RT Master Mix (TaKaRa, Japan). The Power SYBR Green PCR Master Mix (Thermo Fisher, USA) was applied for the quantitative real-time PCR of target mRNA detection using ABI 7300 Fast Real-Time PCR Systems (Applied Biosystem, USA). For miRNA expression study, cDNA were reversely transcribed from RNA samples and detection of miR-378 expression level were conducted with using miRCURY LNA™ Universal RT microRNA PCR kit (Exiqon, Vedbaek, Denmark) following the manufacturer’s manual. The relative fold changes of candidate genes were analyzed by using 2^−ΔΔCt^ method.

### Western blot analysis

Total protein of harvested cell were lysed using RIPA buffer (25 mM Tris-Cl, pH 8.0, 150 mM NaCl, 0.1% SDS, 0.5% sodium deoxycholate, 1% NP-40) supplemented with complete mini protease inhibitor cocktail (Roche, USA) and the soluble protein was collected by centrifuge at 14,000 rpm for 10 min at 4°C. Soluble protein fractures were then mixed with 5x sample loading buffer (Roche, USA) and boiled for 5 min. To perform Western blotting analysis, the protein samples were subjected to SDS-PAGE gel and electrophoresed at 120V for 2h. After that, the protein from SDS-PAGE gel was electroblotted onto a PVDF membrane at 100 V for 1 h at 4°C. The membranes were then blocked with 5% non-fat milk and probed with the following antibodies: β-catenin (1: 3,000, BD Biosciences, USA), Wnt6 (1: 1,000, Sigma, USA), Wnt10a (1: 1,000) and β-actin (1: 4,000, Sigma, USA). The results were visualized on the X-Ray film by Kodak film developer (Fujifilm, Japan).

### Luciferase assay

Dual-luciferase assay was performed according to the instructions of dual-luciferase assay reagent (Promega, USA) with some modifications. Briefly, HEK293 cells were seeded in 24-well plate, and the cells were allowed to grow until 80% confluence. Cells were then transfected with TOPFlash or constructed pmiR-GLO luciferase reporter together with miR-378 mimics or antagonists. The pRSV-β-galactoside (ONPG) vector was co-transfected as normalization control. The plate was placed into a PerkinElmer VictorTM X2 2030 multilateral reader (Waltham, USA) to measure the firefly luciferase activity as well as the β-galactosidase activity. The ratio of firefly luciferase to β-galactosidase activity in each sample was revealed as a measurement of the normalized luciferase activity. All experiments were performed in triplicate.

### Animal surgery and cell local injection

A standard mouse open transverse femoral fracture model was applied in this study.^31^ For bone fracture healing capacity comparison of WT and miR-378 TG mouse, 10 male mice of each type at the age of three month was used. For therapeutic effect of sh-miR-378 lentivirus-infected MSC on bone fracture healing, 20 of miR-378 TG mice at the age of three months were applied. Generally speaking, the mice were carried under general anesthesia and sterilizing procedures. A lateral incision was made through shaved skin from right lateral knee to the greater trochanter, the osteotomy was made by a hand saw at the middle site of right femur, then a hole was drilled at the intercondylar notch by inserting a 23-gauge needle (BD Biosciences, USA) into the bone marrow cavity to fix the fracture. The incision was closed and the fracture was confirmed by X-radiography. For sh-miR-378 treatment study, 20 of miR-378 TG mice were equally and randomly assigned into 2 groups after surgery: sh-NC group and sh-miR-378 group. The mouse MSCs were harvested by 0.25% trypsin and re-suspended in PBS. A total of 5x 10^5^ cells resuspended in 20µl PBS were locally injected into the fracture site of bone under anesthesia using a Hamilton syringe (Hamilton, USA) and a 30-gauge 1/2 needle (BD Biosciences, USA). X-rays were taken weekly using a digital X-ray machine (Faxitron X-Ray Corp., USA) to evaluate the fracture healing condition. Each mouse was exposure for 6000 ms and at voltage of 32 kV. Four weeks post surgery, the mice were sacrificed and the femurs were collected for analysis.

### Micro-computed tomography (μCT)

The structure differences of mouse femur and tibia, as well as the structural change on the fracture sites were quantitatively assessed using μCT as previously described.^32^ Mice femur and tibia samples were excised and the muscles and soft tissues were carefully removed. All the specimens were imaged using a high-solution μCT40 (Scanco Medical, Switzerland) with a voltage of 70kV and a current of 114μA and 10.5μm isotropic resolution. The gray-scale images were segmented with a low-pass filter (sigma=1.2, support=2) to suppress noise and a fixed threshold to perform three dimensional (3D) reconstruction of the mineralized bone phase. Reconstruction of low and high density mineralized bone were performed using different thresholds (low attenuation=158, high attenuation=211) following the previously published protocol with a small modification.^33^ The high-density tissues represented the newly formed highly mineralized bone, while the low ones represented the newly formed callus. Bone volume (BV), tissue volume (TV) and BV/TV of each specimen was recorded for analysis.

### Three-point bending mechanical testing

Mechanical testing was performed within 24h after termination at room temperature. The femurs were loaded on a three-point bending device (H25KS; Hounsfield Test Equipment Ltd., UK) in the anterior-posterior direction with the span of the two blades set as 8 mm. The force loading point was set at the fracture site. The femurs were tested to failure with a constant displacement rate of 6mm/min. Test of contralateral intact femur was also included as an internal control. After test, femur loading displacement curve was generated automatically by built-in software (QMAT Professional Material testing software; Hounsfield Test Equipment Ltd.). The modulus of elasticity (E-modulus) which represents tissue stiffness, ultimate load, and energy to failure which indicates tissue toughness were obtained by previous mentioned software. The biomechanical properties of newly formed bone were further analyzed and expressed as percentages of the contralateral intact bone properties.

### Histology and immunohistochemistry

All femurs and tibia were initially fixed with 10% formalin for 48 h followed by decalcification in 10 % EDTA solution for 2 weeks. The femurs and tibia were then embedded in paraffin and sliced into 5-μm sections by a rotary microtome (HM 355S, Thermo Fisher, USA). For bone morphology study, the femurs and tibia were sliced along their long axis in the coronal plane and short axis in the transverse plane, respectively. For bone fracture study, the femurs were sliced along the long axis in the coronal plane. After deparaffinization, immunohistochemistry (IHC) staining was performed. The sections were stained with hematoxylin & eosin (H&E) for histomorphometric analysis. IHC staining was performed using a standard protocol as previously reported.^31^ Secretions were treated with primary antibodies to OCN (1:100, Abcam, USA) and OPN (1:100, Abcam, USA) overnight at 4°C. The horseradish peroxidase-streptavidin system (Dako, USA) was applied for IHC signal detection, followed by counterstaining with hematoxylin. The images of positive stained cells in the fracture site were captured using light microscope (Leica, Cambridge, UK).

### Statistical analysis

Experimental data are expressed as Mean ± SD. Two groups of data were statistically analyzed using Mann-Whitney U test. The results were considered to be statistically significant when *p* < 0.05.

## Author Contributions

G.L. and J.F.Z. spearheaded and supervised all the experiments. G.L., J.F.Z., D.H.Z. and L.F. designed research. L.F., L.S., Z.M.Y., T.Y.W., H.X.W., W.P.L., and Y.F.L. conducted experiments. L.F. and J.F.Z. analyzed data. D.H.Z. provided materials. L.F., G.L., H.T.L. and J.F.Z. prepared the manuscript. All authors reviewed and approved the manuscript.

## Funding supports

This work was supported by grants from the National Natural Science Foundation of China (81430049, 81772322 and 81772404) and Hong Kong Government Research Grant Council, General Research Fund (14120118, 14160917, 9054014 N_CityU102/15 and T13-402/17-N). This study was also supported in part by SMART program, Lui Che Woo Institute of Innovative Medicine, The Chinese University of Hong Kong and the research was made possible by resources donated by Lui Che Woo Foundation Limited.

---

**Supplementary Figure 1.**
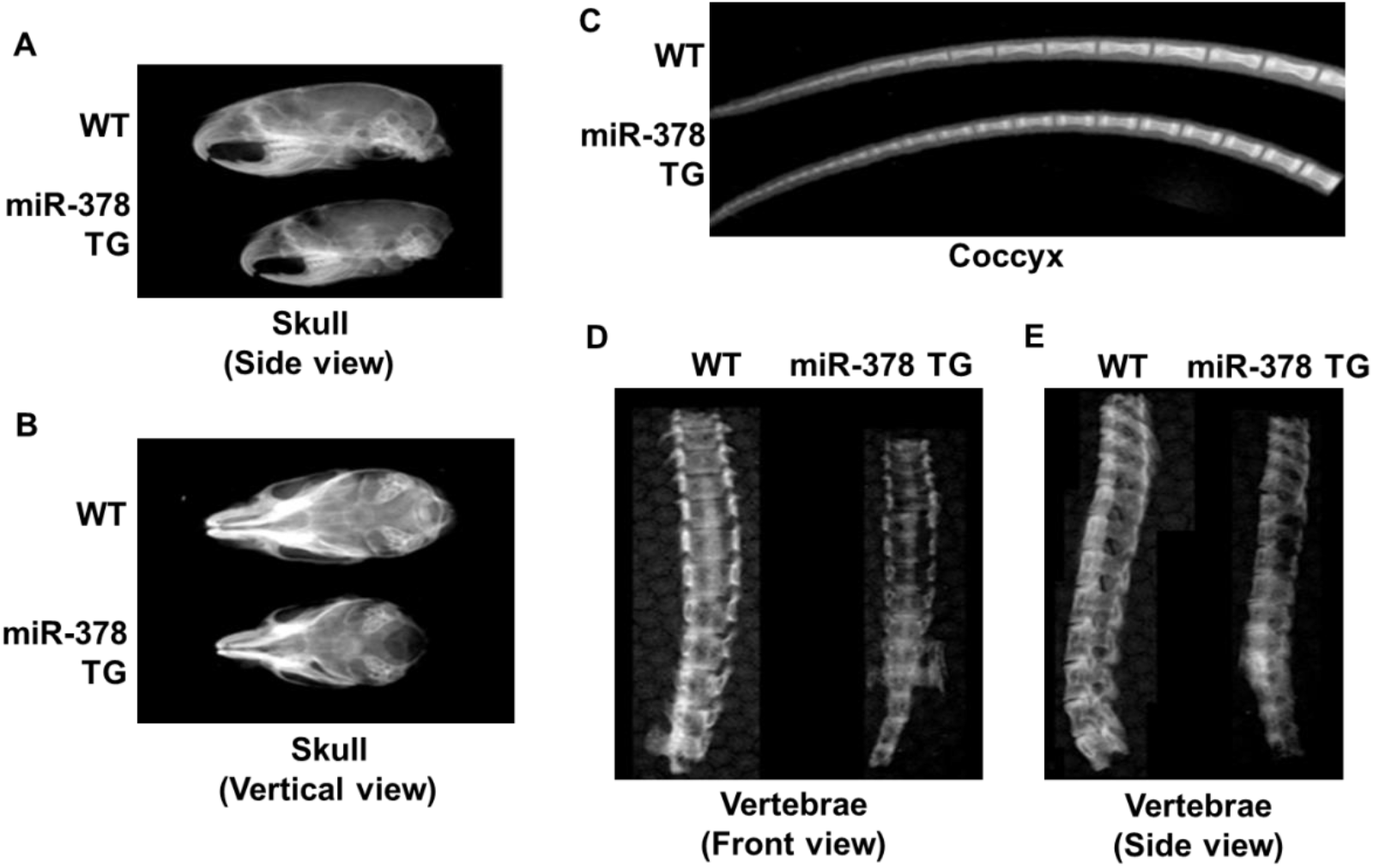
The bone phenotype of skull (**A** for side view and **B** for vertical view), tail **(C)** and spine (**D** for front view and **E** for side view) of miR-378 TG mice and their wild-type (WT) mice were examined by digital radiography.

**Supplementary Figure 2.**
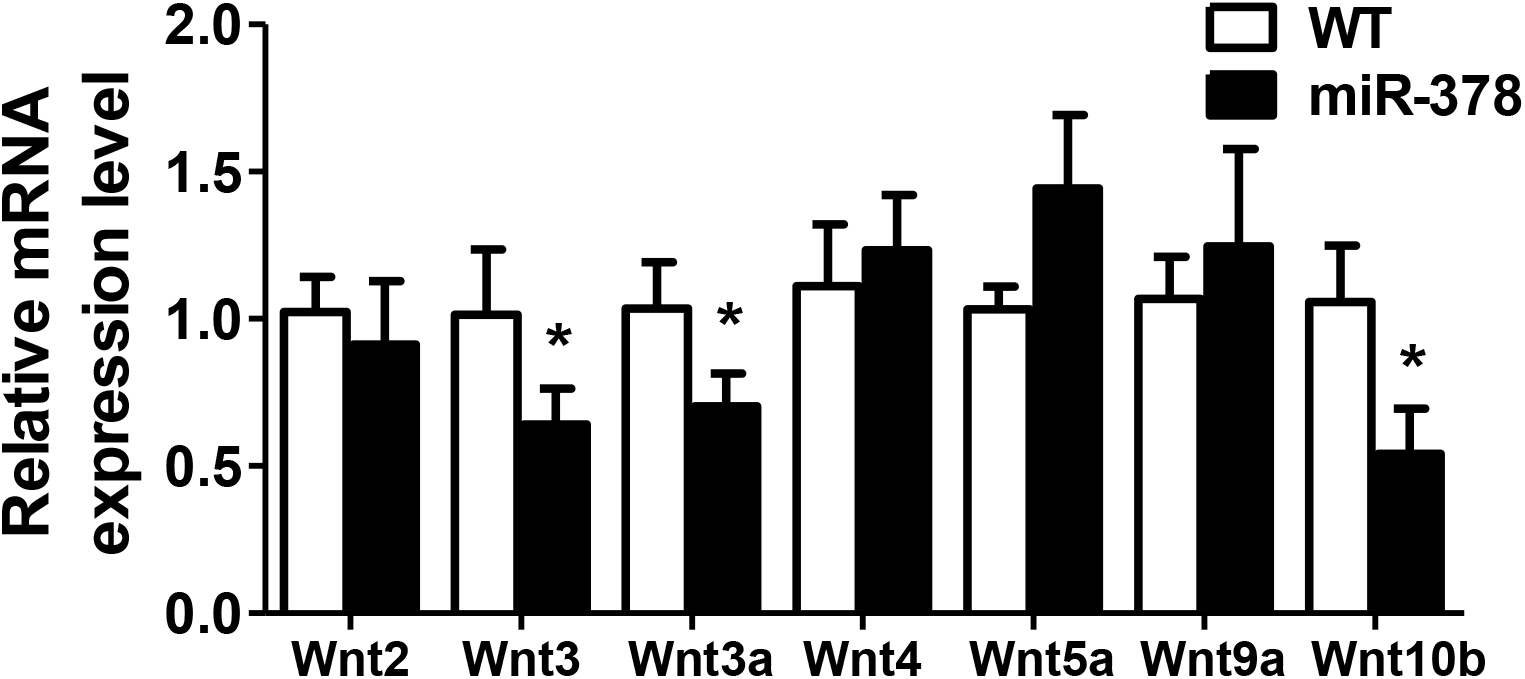
The mRNA expression level of Wnt family members which could activate Wnt/β-catenin signaling pathway revealed by Real-time PCR

**Supplementary Figure 3.**
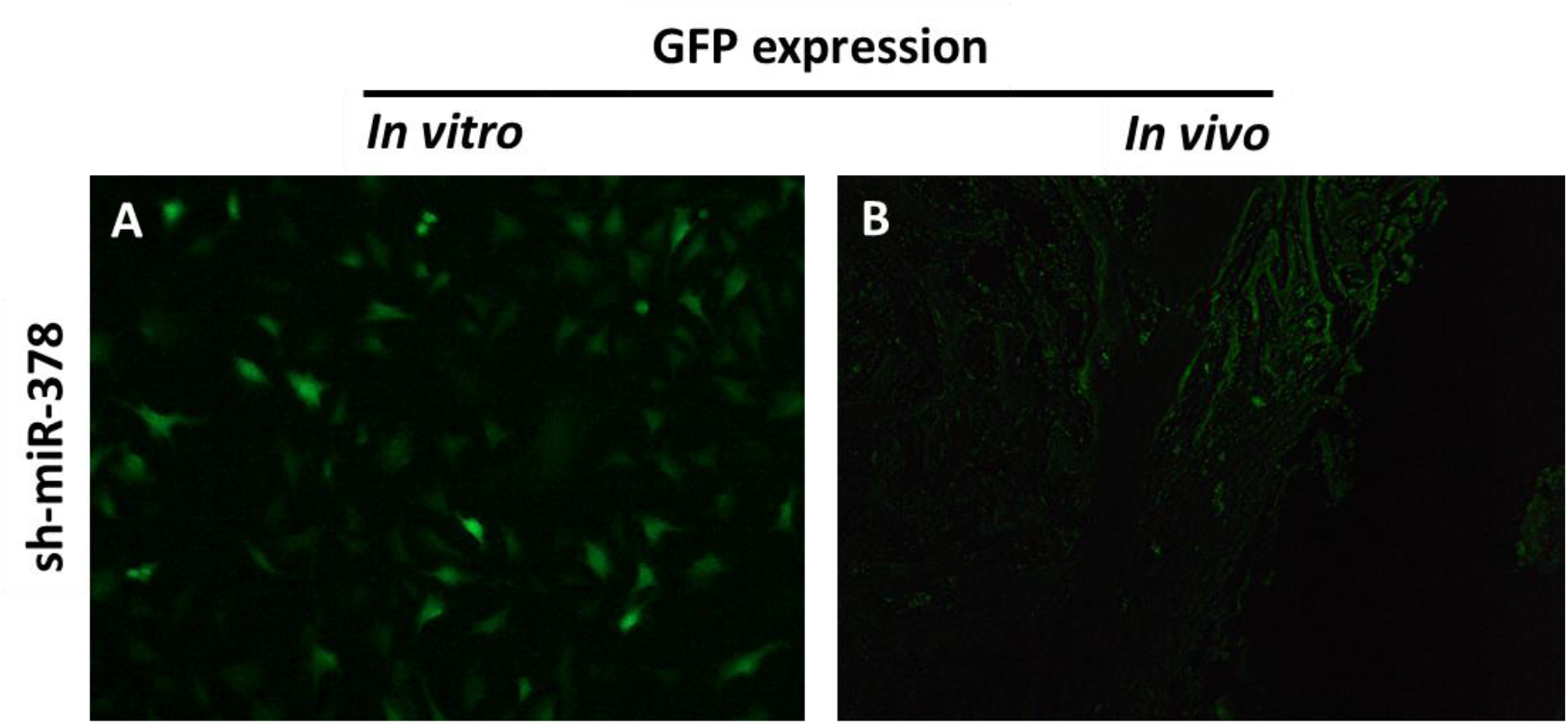
The GFP expression was detected in sh-miR-378 infected BMSCs. **A.** Scramble or sh-miR-378 infected BMSCs *in vitro* (GFP positive cells) **B.** The sh-miR-378 infected MSCs in fracture callus 4 weeks after bone fracture (GFP positive cells).

**Supplementary Figure 4.**
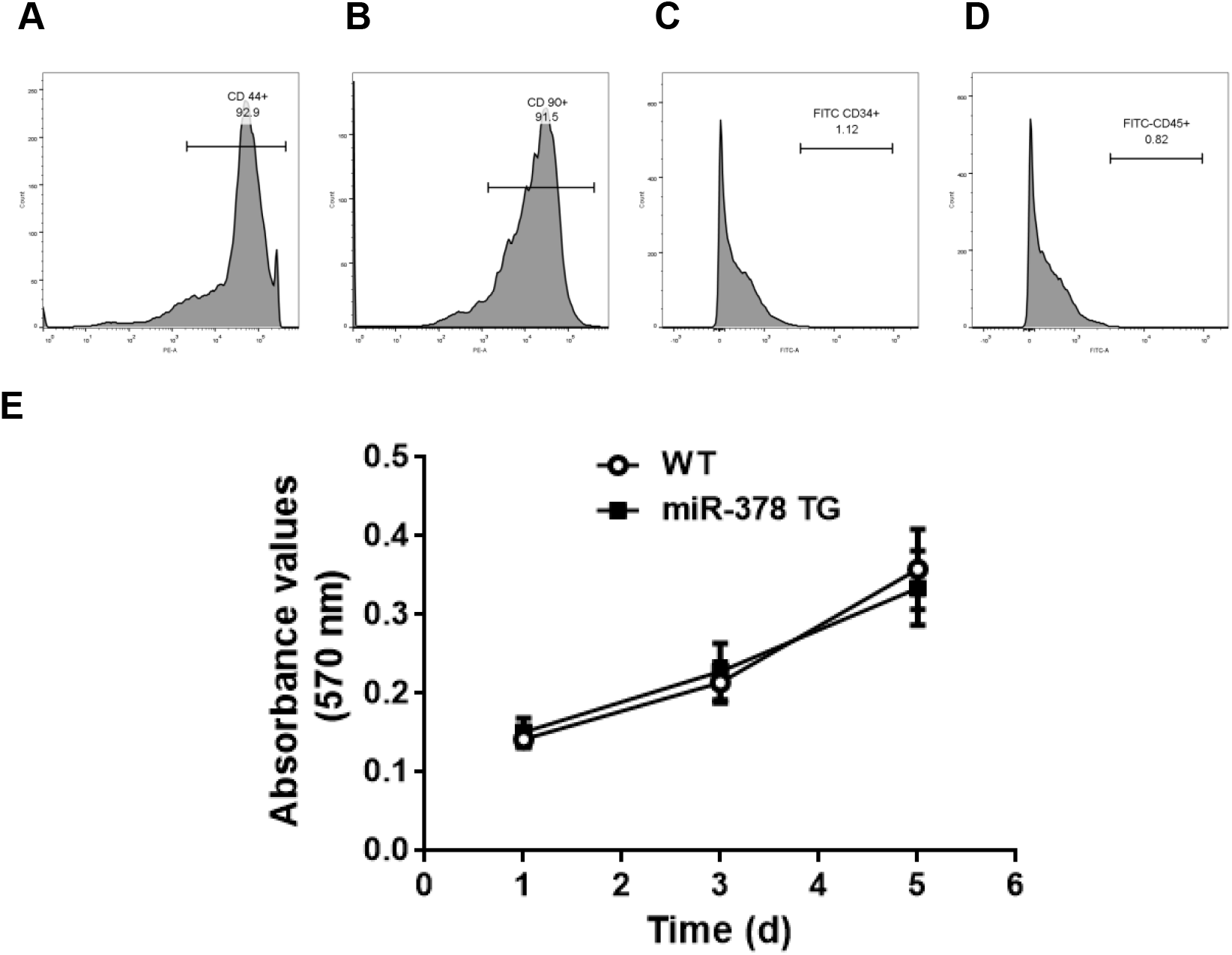
Confirmation of mouse bone marrow MSCs by using flow cytometry and study of MSC proliferation activities. **(A&B)** Flow cytometry analysis results showed that these cells were positive for MSC markers CD44 (A) and CD90 (B). **(C)** Cells were negative for endothelial cell marker CD34. **(D)** Cells were negative for haematopoietic cell marker CD45. **(E)** The MTT assay of MSCs isolated from WT and miR-378 TG mice. The two cells showed no obviously different proliferation activity.

**Supplementary Figure 5.**
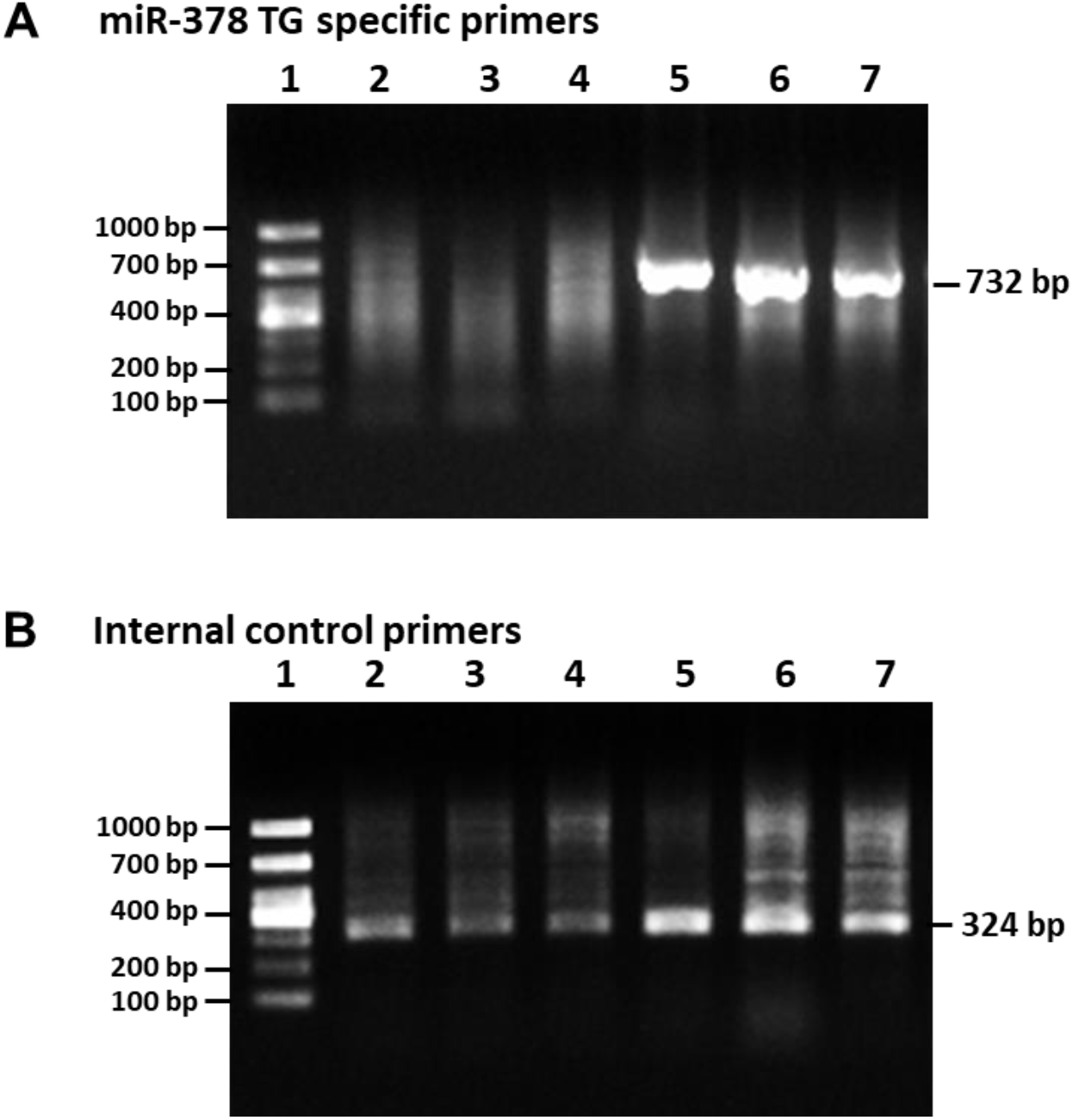
Gel electrophoresis of PCR products using specific primers for the genotyping of miR-378 TG mice. **(A)** miR-378 TG specific primers: 732 bp. **(B)** Internal control primers: 324 bp. Lane 1: 100 bp DNA ladder. Lane 2-4: WT mice genotype. Lane 5-7: miR-378 TG mice genotype.

## Reference

1. Carrer, M, Liu, N, Grueter, CE, Williams, AH, Frisard, MI, Hulver, MW, et al. (2012). Control of mitochondrial metabolism and systemic energy homeostasis by microRNAs 378 and 378*. Proc Natl Acad Sci U S A 109: 15330–15335.

2. Huang, K, Fu, J, Zhou, W, Li, W, Dong, S, Yu, S, et al. (2014). MicroRNA-125b regulates osteogenic differentiation of mesenchymal stem cells by targeting Cbfbeta in vitro. Biochimie 102: 47–55.

3. Xie, Q, Wang, Z, Zhou, H, Yu, Z, Huang, Y, Sun, H, et al. (2016). The role of miR-135-modified adipose-derived mesenchymal stem cells in bone regeneration. Biomaterials 75: 279–294.

4. Li, Z, Hassan, MQ, Volinia, S, van Wijnen, AJ, Stein, JL, Croce, CM, et al. (2008). A microRNA signature for a BMP2-induced osteoblast lineage commitment program. Proc Natl Acad Sci U S A 105: 13906–13911.

5. Kim, YJ, Bae, SW, Yu, SS, Bae, YC, and Jung, JS (2009). miR-196a regulates proliferation and osteogenic differentiation in mesenchymal stem cells derived from human adipose tissue. J Bone Miner Res 24: 816–825.

6. Zhang, JF, Fu, WM, He, ML, Xie, WD, Lv, Q, Wan, G, et al. (2011). MiRNA-20a promotes osteogenic differentiation of human mesenchymal stem cells by co-regulating BMP signaling. RNA Biol 8: 829–838.

7. Zhang, JF, Fu, WM, He, ML, Wang, H, Wang, WM, Yu, SC, et al. (2011). MiR-637 maintains the balance between adipocytes and osteoblasts by directly targeting Osterix. Mol Biol Cell 22: 3955–3961.

8. Hupkes, M, Sotoca, AM, Hendriks, JM, van Zoelen, EJ, and Dechering, KJ (2014). MicroRNA miR-378 promotes BMP2-induced osteogenic differentiation of mesenchymal progenitor cells. BMC Mol Biol 15: 1.

9. You, L, Gu, W, Chen, L, Pan, L, Chen, J, and Peng, Y (2014). MiR-378 overexpression attenuates high glucose-suppressed osteogenic differentiation through targeting CASP3 and activating PI3K/Akt signaling pathway. Int J Clin Exp Pathol 7: 7249–7261.

10. Kahai, S, Lee, SC, Lee, DY, Yang, J, Li, M, Wang, CH, et al. (2009). MicroRNA miR-378 regulates nephronectin expression modulating osteoblast differentiation by targeting GalNT-7. PLoS One 4: e7535.

11. Yu, J, Kong, X, Liu, J, Lv, Y, Sheng, Y, Lv, S, et al. (2014). Expression profiling of PPARgamma-regulated microRNAs in human subcutaneous and visceral adipogenesis in both genders. Endocrinology 155: 2155–2165.

12. Yu, BL, Peng, XH, Zhao, FP, Liu, X, Lu, J, Wang, L, et al. (2014). MicroRNA-378 functions as an onco-miR in nasopharyngeal carcinoma by repressing TOB2 expression. Int J Oncol 44: 1215–1222.

13. Chen, LT, Xu, SD, Xu, H, Zhang, JF, Ning, JF, and Wang, SF (2012). MicroRNA-378 is associated with non-small cell lung cancer brain metastasis by promoting cell migration, invasion and tumor angiogenesis. Med Oncol 29: 1673–1680.

14. Ma, J, Lin, J, Qian, J, Qian, W, Yin, J, Yang, B, et al. (2014). MiR-378 promotes the migration of liver cancer cells by down-regulating Fus expression. Cell Physiol Biochem 34: 2266–2274.

15. Xu, LL, Shi, CM, Xu, GF, Chen, L, Zhu, LL, Zhu, L, et al. (2014). TNF-alpha, IL-6, and leptin increase the expression of miR-378, an adipogenesis-related microRNA in human adipocytes. Cell Biochem Biophys 70: 771–776.

16. Huang, N, Wang, J, Xie, W, Lyu, Q, Wu, J, He, J, et al. (2015). MiR-378a-3p enhances adipogenesis by targeting mitogen-activated protein kinase 1. Biochem Biophys Res Commun 457: 37–42.

17. Liu, SY, Zhang, YY, Gao, Y, Zhang, LJ, Chen, HY, Zhou, Q, et al. (2015). MiR-378 Plays an Important Role in the Differentiation of Bovine Preadipocytes. Cell Physiol Biochem 36: 1552–1562.

18. Zhang, Y, Li, C, Li, H, Song, Y, Zhao, Y, Zhai, L, et al. (2016). miR-378 Activates the Pyruvate-PEP Futile Cycle and Enhances Lipolysis to Ameliorate Obesity in Mice. EBioMedicine 5: 93–104.

19. Ell, B, Mercatali, L, Ibrahim, T, Campbell, N, Schwarzenbach, H, Pantel, K, et al. (2013). Tumor-induced osteoclast miRNA changes as regulators and biomarkers of osteolytic bone metastasis. Cancer Cell 24: 542–556.

20. Duan, P, and Bonewald, LF (2016). The role of the wnt/beta-catenin signaling pathway in formation and maintenance of bone and teeth. Int J Biochem Cell Biol 77: 23–29.

21. Huang, Y, Liu, X, and Wang, Y (2015). MicroRNA-378 regulates neural stem cell proliferation and differentiation in vitro by modulating Tailless expression. Biochem Biophys Res Commun 466: 214–220.

22. Zeng, M, Zhu, L, Li, L, and Kang, C (2017). miR-378 suppresses the proliferation, migration and invasion of colon cancer cells by inhibiting SDAD1. Cell Mol Biol Lett 22: 12.

23. Yu, F, Fan, X, Chen, B, Dong, P, and Zheng, J (2016). Activation of Hepatic Stellate Cells is Inhibited by microRNA-378a-3p via Wnt10a. Cell Physiol Biochem 39: 2409–2420.

24. Cawthorn, WP, Bree, AJ, Yao, Y, Du, B, Hemati, N, Martinez-Santibanez, G, et al. (2012). Wnt6, Wnt10a and Wnt10b inhibit adipogenesis and stimulate osteoblastogenesis through a beta-catenin-dependent mechanism. Bone 50: 477–489.

25. Tang, N, Song, WX, Luo, J, Luo, X, Chen, J, Sharff, KA, et al. (2009). BMP-9-induced osteogenic differentiation of mesenchymal progenitors requires functional canonical Wnt/beta-catenin signalling. J Cell Mol Med 13: 2448–2464.

26. Gaur, T, Lengner, CJ, Hovhannisyan, H, Bhat, RA, Bodine, PV, Komm, BS, et al. (2005). Canonical WNT signaling promotes osteogenesis by directly stimulating Runx2 gene expression. J Biol Chem 280: 33132–33140.

27. Semenov, MV, and He, X (2006). LRP5 mutations linked to high bone mass diseases cause reduced LRP5 binding and inhibition by SOST. J Biol Chem 281: 38276–38284.

28. James, MJ, Jarvinen, E, Wang, XP, and Thesleff, I (2006). Different roles of Runx2 during early neural crest-derived bone and tooth development. J Bone Miner Res 21: 1034–1044.

29. Jing, H, Liao, L, An, Y, Su, X, Liu, S, Shuai, Y, et al. (2016). Suppression of EZH2 Prevents the Shift of Osteoporotic MSC Fate to Adipocyte and Enhances Bone Formation During Osteoporosis. Mol Ther 24: 217–229.

30. Huang, S, Xu, L, Sun, Y, Wu, T, Wang, K, and Li, G (2015). An improved protocol for isolation and culture of mesenchymal stem cells from mouse bone marrow. J Orthop Translat 3: 26–33.

31. Sun, Y, Xu, L, Huang, S, Hou, Y, Liu, Y, Chan, KM, et al. (2015). mir-21 overexpressing mesenchymal stem cells accelerate fracture healing in a rat closed femur fracture model. BioMed research international 2015: 412327.

32. Hauser, S, Wulfken, LM, Holdenrieder, S, Moritz, R, Ohlmann, CH, Jung, V, et al. (2012). Analysis of serum microRNAs (miR-26a-2*, miR-191, miR-337-3p and miR-378) as potential biomarkers in renal cell carcinoma. Cancer Epidemiol 36: 391–394.

33. Xu, J, Wang, B, Sun, Y, Wu, T, Liu, Y, Zhang, J, et al. (2016). Human fetal mesenchymal stem cell secretome enhances bone consolidation in distraction osteogenesis. Stem Cell Res Ther 7: 134.

